# SYS-1/beta-catenin inheritance and regulation by Wnt-signaling during asymmetric cell division

**DOI:** 10.1101/2023.07.21.550069

**Authors:** Maria F. Valdes Michel, Bryan T. Phillips

## Abstract

Asymmetric cell division (ACD) allows daughter cells of a polarized mother to acquire different developmental fates. In *C. elegans*, the Wnt/β-catenin Asymmetry (WβA) pathway oversees many embryonic and larval ACDs; here, a Wnt gradient induces an asymmetric distribution of Wnt signaling components within the dividing mother cell. One terminal nuclear effector of the WβA pathway is the transcriptional activator SYS-1/β-catenin. SYS-1 is sequentially negatively regulated during ACD; first by centrosomal regulation and subsequent proteasomal degradation and second by asymmetric activity of the β-catenin “destruction complex” in one of the two daughter cells, which decreases SYS-1 levels in the absence of WβA signaling. However, the extent to which mother cell SYS-1 influences cell fate decisions of the daughters is unknown. Here, we quantify inherited SYS-1 in the differentiating daughter cells and the role of SYS-1 inheritance in Wnt-directed ACD. Photobleaching experiments demonstrate the GFP::SYS-1 present in daughter cell nuclei is comprised of inherited and *de novo* translated SYS-1 pools. We used a photoconvertible DENDRA2::SYS-1, to directly observe the dynamics of inherited SYS-1. Photoconversion during mitosis reveals that SYS-1 clearance at the centrosome preferentially degrades older SYS-1, and this accumulation is regulated via dynein trafficking. Photoconversion of the EMS cell during Wnt-driven ACD shows daughter cell inheritance of mother cell SYS-1. Additionally, loss of centrosomal SYS-1 increased inherited SYS-1 and, surprisingly, loss of centrosomal SYS-1 also resulted in increased levels of *de novo* SYS-1 in both EMS daughter cells. Lastly, we show that daughter cell negative regulation of SYS-1 via the destruction complex member APR-1/APC is key to limit both the *de novo* and the inherited SYS-1 pools in both the E and the MS cells. We conclude that regulation of both inherited and newly translated SYS-1 via centrosomal processing in the mother cell and daughter cell regulation via Wnt signaling are critical to maintain sister SYS-1 asymmetry during ACD.

## Introduction

Asymmetric cell division (ACD) controls cell fate specification during development by increasing cell diversity in developing tissues, including such diverse examples as *C. elegans* embryos*, Drosophila* neuroblasts, and mammalian lung epithelial cells [1–4]. During adulthood, ACD maintains tissue integrity by promoting stem cell self-renewal while producing cells that will further differentiate to repopulate dying or degraded tissues [5]. Given that ACDs of stem cells maintain the ability to replenish damaged or growing tissues even post-development, tight regulation of these stem cell niches is needed to prevent tumorigenesis. Accordingly, pathways that regulate ACD are implicated in the increased risk of cancer [6]. The same signaling pathways often regulate of both homeostatic and developmental ACDs, thus elucidating the mechanisms regulating ACD will contribute to the understanding of cell fate specification during development and illuminate the mechanism by which tumors develop [7].

The Wnt/β-catenin pathway is a well-conserved signaling pathway that regulates stem cell maintenance, cell renewal and differentiation, cell polarity, and ACD [8–11]. In this pathway, Wnt ligands activate a signal transduction cascade that stabilizes the coactivator β-catenin, which translocates into the nucleus, binds TCF transcription factors and initiates transcription of Wnt target genes. In the absence of Wnt ligand, β-catenin is degraded by a “destruction complex” comprised of casein kinase 1α (CK1α), glycogen synthase kinase 3β (GSK3β), and two scaffolds, Axin and adenomatous polyposis coli (APC). In the absence of β-catenin, TCF acts as a transcriptional repressor of Wnt-target genes [12–14]. Aberrant regulation of Wnt signaling leads to developmental cell specification defects and is a common theme of cancer biology. Tumor sequencing further reveals frequent inactivating mutations in APC and activating mutations in β-catenin [15–17]. Constitutive Wnt pathway activity is present in 90% of colorectal cancer cases. This may be impacted by changes in the cell polarity and in ACD, which leads to an increase in the cellular proliferation of intestinal epithelial stem cells [16, 18]. Though regulation of β-catenin is essential for proper Wnt signaling, how the mother cell distributes β-catenin into the daughter cells and how β-catenin is differentially regulated during ACD remains unclear.

The Wnt/β-catenin asymmetry (WβA) pathway in *C. elegans* controls many embryonic and larval ACDs through regulation of the β-catenin, SYS-1 [19–21]. The WβA pathway is a branched pathway specialized for ACD that uses a β-catenin paralog, WRM-1 and SYS-1, in each branch [19, 22–26]. The WRM-1 branch activates the nuclear export of approximately 50% of the TCF/LEF DNA-binding protein, POP-1, via association with WRM-1 [22]. WRM-1-assisted export of POP-1 is, counterintuitively, required for Wnt-target gene activation because POP-1 functions as a transcriptional repressor when unbound to β-catenin/SYS-1 [22, 25]. POP-1 nuclear export enables the remaining nuclear POP-1 to complex with its coactivator β-catenin/SYS-1 [19, 27]. The second branch of the WβA pathway regulates the stabilization and subsequent nuclear localization of SYS-1, which activates transcription of Wnt-target genes by complexing with POP-1 [28, 29].

SYS-1 is broadly expressed at the transcriptional level but expression, including asymmetric expression, is tightly regulated at the protein level [21, 23]. One example of regulation of SYS-1 protein expression is via centrosomal localization and subsequent turnover. SYS-1 symmetrically localizes to the centrosomes during ACD, and that SYS-1 centrosomal localization is dependent on the centrosomal protein, RSA-2 [21, 23, 30–33]. After ACD in the absence of RSA-2, overall SYS-1 protein levels, including nuclear SYS-1, increase in both daughter cells [30, 33]. These data suggest centrosomal regulation leads to the negative regulation and subsequent degradation of centrosomally localized SYS-1 and limits SYS-1 levels in daughter cells. Depletion of the proteasome core ATPase subunit of the 26S proteasome, RPT-4, results in elevated and stabilized centrosomal SYS-1 and decreased SYS-1 turnover compared to wild type [30, 33]. These results are consistent with the hypothesis that the centrosomal proteasome plays a key role in regulating SYS-1 turnover at the centrosome via active degradation by the proteasome.

The microtubule motor dynein enhances the rapid accumulation of SYS-1 on mitotic centrosomes of large embryonic blastomeres via microtubule trafficking of SYS-1 [34], leading to SYS-1 mitotic turnover. Thus, dynein works as a negative regulator of SYS-1, by enhancing centrosomal SYS-1 levels and subsequent degradation by the centrosomal proteasome. Knockdown of single dynein subunits, including heavy chain DHC-1 or the light chain subunits DLC-1 and DYLT-1, lead to reduced SYS-1 centrosomal enrichment and elevated SYS-1 in both daughter cells. Additionally, loss of these dynein complex subunits results in the conversion of larval ACDs to symmetric divisions, increasing the Wnt-signaled cell fates [34]. These data highlight the importance of microtubule trafficking for SYS-1 centrosomal localization and subsequent degradation. However, we previously showed that, when the worm homolog of mammalian proteasomal trafficking factor ECM29 [35], *ecps-1,* is depleted, embryos with increased centrosomal SYS-1 were observed, in addition to a second substrate of the centrosomal proteasome, ZYG-1 [34, 36]. Thus, the current model suggests that SYS-1 and the proteasome are both actively trafficked via dynein to the centrosome during mitosis and, upon proper localization, result in SYS-1 degradation.

While the above model details SYS-1 negative regulation via centrosomal localization during mitosis in the mother cell, SYS-1 is also regulated after an ACD by the β-catenin destruction complex under the regulatory control of WβA in the daughter cells. In response to the Wnt ligand gradient, the positive Wnt regulators, Frizzled and Disheveled, are asymmetrically localized at the posterior cortex, resulting in their enrichment in Wnt-signaled daughter cell after cytokinesis [37–40]. Members of the destruction complex, APC and Axin, are localized at the anterior pole of the mother cell and are asymmetrically enriched in the anterior Wnt-unsignaled daughter cell. The worm homolog of APC, APR-1, is essential for SYS-1 negative regulation during ACD [29, 41, 42]. These data, coupled with the above centrosomal regulation data, suggest the current model of SYS-1 regulation occurs via centrosomal degradation of mother cell SYS-1 and asymmetric regulation of *de novo* synthesized SYS-1 by the destruction complex in the Wnt-signaled and -unsignaled daughter cells [28, 29].

Lineage studies show that the Wnt-signaled daughters of Wnt-signaled cells respond more strongly to Wnt signaling in the subsequent division compared to their cousins, the Wnt-signaled daughters of Wnt-unsignaled cells [43, 44]. The heightened response can be visualized by both an enrichment of SYS-1 in the nucleus and increased target gene transcription [43, 44]. This nuclear SYS-1 enrichment suggests SYS-1 may be shielded from proteasomal degradation during nuclear localization, allowing for greater SYS-1 inheritance into the daughter cells of previously Wnt-signaled mother cells and progressively enriching the Wnt-signaled lineages for SYS-1 over time. Alternatively, enrichment of positive regulators of SYS-1 in Wnt-signaled lineages, such as Frizzled or Dishevelled [45, 46], would also be predicted to increase downstream SYS-1 stabilization. In this latter case, inheritance of upstream Wnt pathway components would be expected to similarly increase WRM-1 and decrease nuclear POP-1 levels between cousins, which was indeed observed in this study [12, 43, 44]. This suggests that increased SYS-1 inheritance from signaled mothers is unlikely to be the sole mechanism of cousin enrichment. However, the extent to which direct SYS-1 inheritance contributes to a Wnt pathway “memory” during normal ACD and how Wnt signaling regulates inherited and *de novo* SYS-1 remains unknown.

Here, we show centrosomal degradation of SYS-1 in the EMS mother cell is incomplete, leading to SYS-1 inheritance in the E and the MS daughter cells during wild-type ACDs. We developed a photoconvertible SYS-1 to observe the dynamics of preexisting and newly synthesized SYS-1 at the centrosome during mitosis. We found that preexisting and newly synthesized SYS-1 levels are trafficked to the mitotic centrosome via the dynein complex where they are both degraded. SYS-1 coupling, trafficking, and degradation at the mother cell centrosome are necessary for maintaining sister cell asymmetry and wild-type SYS-1 asymmetry in E and MS daughter cells during ACD. Surprisingly, *de novo* SYS-1 increases in both daughter cells after the over-inheritance of mother cell SYS-1 during asymmetric cell division. Additionally, depletion of the negative regulator APR-1 increased both pools of SYS-1 in the E and MS nuclei, suggesting that, when inherited SYS-1 is affected, *de novo* SYS-1 is also misregulated and vice versa. Thus, we propose a model in which SYS-1 centrosomal degradation is a key regulator of SYS-1 inheritance that prevents the loss of daughter cell asymmetry. Further regulation of inherited and *de novo* SYS-1 in the E and MS daughter cells via Wnt signaling supports proper sister cell asymmetry due to a free exchange between the inherited and *de novo* SYS-1 pools.

## Methods

### Strains

Strains were maintained on OP50-inoculated nematode growth media (NGM) plates at 20°C (PHX5708 and N2) or 25°C (TX964) using typical *C. elegans* methods [47]. The genotypes of the investigated are as follows: N2 (wild type); TX964 (unc-119(ed3) III; him-3(e1147) IV; teIs98 [P_pie-1_::GFP::SYS-1]); BTP254 *(syb5708[*P_sys-1::_DENDRA2::SYS-1]).

### CRISPR/Cas9 mediated fluorescent tagging of endogenous SYS-1

An in-frame N-terminal DENDRA2-GAS tag was inserted in the endogenous *sys-1* gene prior to the endogenous start codon (SunyBioTech, Fujian, China). Two synonymous nucleotide changes were generated in the donor sequence to prevent the donor from being recognized and cut by Cas9. Genome edited animals were sequence verified with DENDRA-2-*sys-1* specific primers and backcrossed to wild-type N2 worms to reduce the chances of extraneous genetic variation or off target insertions impacting our experiments. The resulting strain, *sys-1(syb5708[*P_sys-1::_DENDRA2::SYS-1]), is designated by strain name BTP254.

### RNAi

To perform RNAi knockdown of target genes, we used HT115 bacteria containing the pL4440 plasmid with a T7-flanked target gene insert. *rsa-2*, *apr-1* and *ecps-1* were obtained from a library supplied by Ahringer (Addgene) [48]. *dlc-1* was obtained from the Phillips lab library and made as described in Thompson *et* al., 2022. Briefly, *dlc-1* was amplified using target gene-flanking primers and *C*.

*elegans* cDNA to generate a new insert. Bacteria with the desired insert were seeded onto isopropyl β-d-1-thiogalactopyranoside (IPTG)-containing plates to induce transcription from the T7 promotor as described previously [49]. Worms were plated onto RNAi bacterial lawns after sodium hypochlorite synchronization as L1s, with the exception of worms plated on *dlc-1* bacterial lawns which were washed from OP50 plates at 24 h before imaging (L3/L4) and plated on to the RNAi plate to avoid larval phenotypes [50, 51].

### Confocal microscopy and FRAP

Confocal microscopy was performed with a Leica SP8 HyD detector system. The objective used was a 63× HC PL APO CS2 objective with N.A. = 1.4, using type F immersion oil. Each analyzed image consisted of 35 summed z-images of the embryo, with each slice approximately 0.6-μm thick. FRAP was performed on the same system. Each embryo was imaged once then photobleached, at anaphase, telophase or cytokinesis. The entire EMS cell was selected as the region of interest (ROI) and bleached at an intermediate focal plane with 100% laser power for 200 iterations. Images were taken immediately after photobleaching at 4, 6, 8 and 10 minutes. Fluorescence intensity was measured on the sum of 6 Z-projection slices in FIJI using the mean fluorescence intensity of ROIs around the nuclei of the E and MS cells.

### Confocal microscopy and photoconversion of DENDRA2::SYS-1

Confocal microscopy was performed via a Leica SP8 HyD detector system using a 63× HC PL APO CS2 objective with N.A. = 1.4 and type F immersion oil. Each image analyzed consisted of the sum of 35 z-images, approximately 0.6-μm thick, across the embryo. Photoconversion of DENDRA2::SYS-1 was performed on the same system. For centrosome and cytoplasm photoconversion the FRAP software was used to photoconvert the desired locale. Before photoconversion, one image was taken in early metaphase to observe and quantify pre-conversion green SYS-1 levels at the centrosome. The ROI was then photoconverted using the UV laser (405nm diode laser) at an intermediate focal plane with 100% laser power for 200 iterations. Excitation light at 488 nm (490–550 nm) or 552 nm (580–670 nm) was provided by HyD lasers set at 30% laser power for both channels on standard mode. Images were taken immediately after photoconversion every 20 seconds through mitosis of the P1 cell with a frame and line average of 2. To obtain the CEI, the mean intensity of the embryonic cytoplasm was subtracted from the centrosomal mean [34].

For whole embryo photoconversion and quantification of DENDRA2::SYS-1 nuclear levels in the E and MS daughter cells, embryos were exposed to a UV laser (405 nm diode laser) at 100% laser power for 36 iterations (frame average: 6, line average: 6) using the normal imaging setting on the Leica SP8 HyD detector system. An image was taken of each embryo prior to photoconversion with the excitation light of 488 nm (490–550 nm) or 552 nm (580-670) HyD lasers set at 30% laser power for both channels on standard mode. Photoconversion was performed at the time of cytokinesis of the EMS cell, and 2 minutes after cytokinesis the timecourse was started, with 1 minute between each image taken for 7 minutes. DENDRA2::SYS-1 nuclear levels were normalized by N2 values utilizing the same settings described above.

## Results

### Photobleaching reveals *de novo* and inherited SYS-1 pools in the E and MS cells and centrosomal SYS-1 dynamics over time

Alterations in centrosomal SYS-1 enrichment correlate with nuclear SYS-1 levels in WβA daughter cells [30, 34]. Disruption of SYS-1 centrosomal levels by knockdown of the centrosomal scaffold protein RSA-2, different dynein subunits (DHC-1, DLC-1, DYLT-1), or the proteasome trafficking adaptor protein ECPS-1 are sufficient to induce extra Wnt-signaled cell fates [34]. These data suggest that impairing mother cell SYS-1 centrosomal localization or degradation increases SYS-1 inheritance in the daughter cells. However, it remains possible that degradation of SYS-1 in the mother cell is also incomplete in wild type and that inherited SYS-1 contributes to normal Wnt-dependent asymmetric cell fate specification. To better understand the dynamics of SYS-1 inheritance in wild type, we used fluorescence recovery after photobleaching (FRAP) in the E and MS daughter cells at different time points. The EMS cell is polarized by Wnt ligand produced in the posterior P2 cell, resulting in E and MS daughter cells with high and low nuclear levels of SYS-1, respectively [24, 39, 52]. EMS was photobleached at different times during mitosis, and nuclear SYS-1 levels in the E and MS daughter cells were measured four minutes after cytokinesis (Fig. 1A,B). If SYS-1 is cleared by the mother cell centrosomal processing, we expect no change between SYS-1 nuclear levels due to EMS photobleaching at different timepoints. Alternatively, if the mother cell SYS-1 protein is inherited by the daughter cells, we expect the largest decrease in SYS-1 levels to occur following a photobleach at cytokinesis. Anaphase photobleach gives slight, non-significant decrease (Fig. 1B), but we speculated that this was due to late SYS-1 synthesis, giving rise to an unbleached population of mother cell SYS-1. To address this, we photobleached at telophase and at cytokinesis. Indeed, after the EMS telophase photobleach, we observed significant decreases in E and MS nuclear SYS-1, suggesting degradation of SYS-1 is incomplete in EMS mother cells (Fig. 1B, S1). Finally, cytokinesis photobleaching, despite the late timing, resulted in visible GFP::SYS-1 in both daughter cell nuclei, revealing both a mother cell contribution and a *de novo* synthesized SYS-1 pool in the E and the MS daughter cells (Fig. 1B). Regardless of the time of photobleaching, the asymmetry between E and MS SYS-1 nuclear levels is maintained (Fig. 1B). These results suggest SYS-1 mother cell clearance is incomplete and that *de novo* SYS-1 is asymmetrically regulated during EMS ACD.

**Figure 1:**
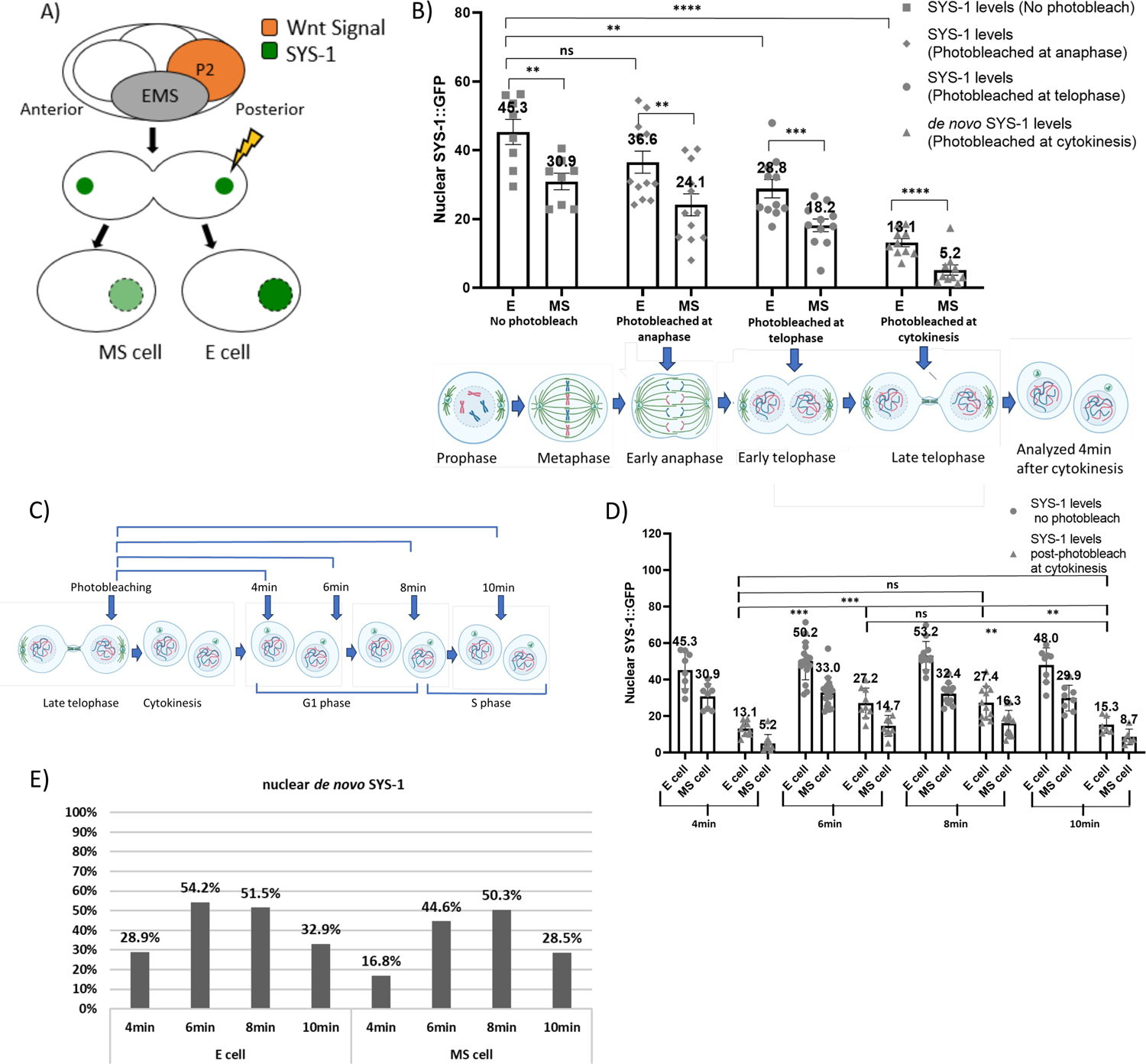
Photobleaching reveals *de novo* and inherited SYS-1 pools in the E and MS cells and centrosomal SYS-1 dynamics over time. A) Diagram of EMS cell FRAP assay. EMS cell is polarized by Wnt from the P2 cell. EMS mother cell is photobleached and SYS-1 nuclear levels are subsequently measured in the E and the MS daughters. B) Quantification of nuclear SYS-1::GFP in E and MS cells without photobleaching (n=8) and with photobleaching at anaphase (n=12), telophase (n=11), and cytokinesis (n=10). Fluorescence recordings were measured 4 min after mother cell division is complete. Mean +/− SEM. Student’s two-tail t-test. C) Photobleaching protocol for panel D) and E). D)Quantification of nuclear fluorescent SYS-1::GFP in the E and MS cells over time in non-bleached controls and EMS cell photobleached at cytokinesis. Mean +/− SEM. Student’s two-tail t-test. E) Percentage of nuclear SYS-1 in daughter cells during a 2 min interval timecourse. N= denoted by grey figures in graph. * p ≤ 0.05, ** p ≤ 0.01, *** p ≤ 0.001, **** p ≤ 0.0001

To observe the dynamics of the *de novo* SYS-1 pool in the daughter cells, we photobleached the EMS mother cell at cytokinesis and measured *de novo* and total nuclear SYS-1 at different times (4, 6, 8, and 10 mins after cytokinesis) (Fig. 1C) with and without photobleaching. Because *de novo* nuclear SYS-1 levels were reduced by 45–83% relative to total SYS-1 levels at each time point, our results confirm SYS-1 inheritance occurs in wild type, however, interestingly, the accumulation of *de novo* SYS-1 was not linear. Instead, we observed a significant increase in nuclear *de novo* SYS-1 levels in both daughter cells during minutes four through six, as expected as total newly synthesized SYS-1 increases (Fig. 1D). However, *de novo* SYS-1 levels plateau around 8 min and then decrease at 10 min, after cytokinesis of the mother cell (Fig. 1D), suggesting *de novo* SYS-1 accumulation ceases or slows throughout the cell cycle. To quantify this change, we calculated the ratio of total to *de novo* nuclear SYS-1 levels over the timecourse in both daughter cells. In the E cell at four minutes, the *de novo* SYS-1 pool represents 28.9% of the total SYS level (Fig. 1E) and increases to 54.2% and 51.5% by minutes 6–8, respectively, after mother cell division. Ten minutes after cytokinesis *de novo* SYS-1 decreases to 33%. Intriguingly, we observe a similar pattern in the MS cell (4 min=17%; 6 min=45%; 8 min=50%; 10 min=29%), though the absolute levels are higher in the Wnt-signaled E daughter (Fig. 1D). This timecourse revealed variable rates of *de novo* SYS-1 synthesis and regulation in the nuclei of the E and MS daughter cells through the cell cycle, suggesting the role of Wnt signaling in SYS-1 stabilization is mostly complete by 8 minutes post-EMS division and that the “unsignaled” MS cell also stabilizes SYS-1, albeit at a lower level than E.

### Photoconversion of DENDRA2::SYS-1 reveals turnover at the centrosome during mitosis

To visualize *de novo* and inherited SYS-1, we generated a photoconvertible *sys-1* transgene, *P_sys–1_*::DENDRA2::SYS-1 that differentiates preexisting (red) and newly synthesized (green) SYS-1 centrosomal accumulation in the P1 cell, which has been previously utilized for SYS-1 centrosomal localization and processing studies [30, 34]. While observing P1 cellular division, in the green channel DENDRA2::SYS-1 is visible at the centrosome prior to photoconversion, consistent with previous studies [30, 34] (Fig. 2A), while the red channel detection shows background levels of centrosomal DENDRA2::SYS-1 (Fig. 2B). To photoconvert DENDRA2::SYS-1 from green to red fluorescence, we exposed the entire P1 cell to a 405nm UV diode laser [53] at 100% power for 36 iterations. After photoconversion, centrosomal DENDRA2::SYS-1 in the green channel sharply decreases (Fig. 2C) and we clearly detect red fluorescent DENDRA2::SYS-1 at the centrosome (Fig. 2D). After confirming successful photoconversion, we measured centrosomal DENDRA2::SYS-1 levels in 20-second increments through P1 mitosis (Fig. 2E). In order to control possible photoconversion by the 488nm laser [54], we followed the same imaging protocol without 405nm photoconversion and followed both emissions over the timecourse. The data were normalized using N2 animals subjected to the same protocol to account for possible photobleaching or changes in development of the embryo induced by the imaging. Our data shows that there is no photoconversion induced by repetitive imaging with the 488nm laser as no changes are observed in the red channel (Fig. S2A). The raw data revealed the preexisting, red centrosomal DENDRA2::SYS-1 decreased over time, likely due to centrosomal degradation. Interestingly, the newly synthesized, green centrosomal DENDRA2::SYS-1 decreased at a slower rate throughout the rest of P1 mitosis (Fig. 2F). Additionally, to quantify photoconversion efficacy, we compared the level of post-conversion red signal to the green signal after mock conversion, both normalized for green pre-conversion signal (Fig. S2A, B). The results indicate ratios of approximately one and no significant difference between the two ratios, indicating successful photoconversion. Together these data suggest that the rate of synthesis and trafficking is relatively equal to the rate of degradation of *de novo* SYS-1 at the centrosome, which is consistent with the previously published idea of a steady-state level of centrosomal GFP::SYS-1 that is stable during the P1 cell mitosis (Vora and Phillips, 2015).

**Figure 2:**
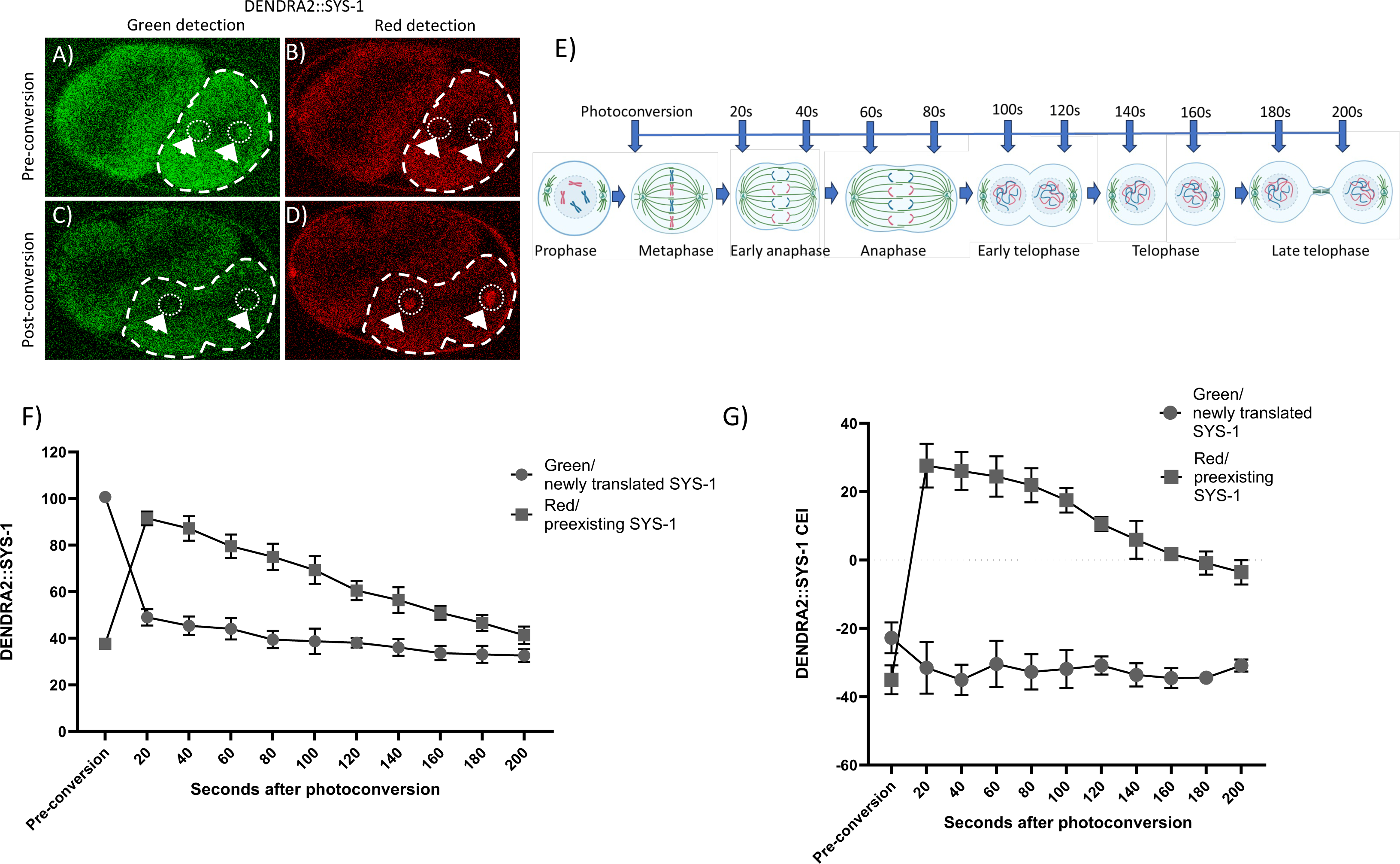
Photoconversion of DENDRA2::SYS-1 reveals turnover at the centrosome during mitosis. A) Pre-conversion image shows centrosomal localization of DENDRA2::SYS-1 in the green channel. B) Centrosomal DENDRA2::SYS-1 is undetectable in the red channel prior to photoconversion by 405nm blue laser exposure. C) After photoconversion newly translated DENDRA2::SYS-1 is trafficked to the centrosome, and D) shows photoconverted DENDRA2::SYS-1 at the centrosome. E) Photoconversion protocol for centrosomal DENDRA2::SYS-1 timecourse. F) Quantification of average centrosomal DENDRA2::SYS-1 in both channels over time. Mean +/− SEM. N=3. G) Centrosomal enrichment index (CEI) of the time course shown in (F). Mean +/− SEM. N=3. CEI was calculated in FIJI by subtracting the mean pixel intensity of a cytoplasmic ROI encompassing the posterior half of the cell with the dotted white line (A–D), from a circular centrosome ROI.

To specifically evaluate the enrichment of DENDRA2::SYS-1 at the centrosome, we used the centrosomal enrichment index (CEI) [34]. CEI measures the degree to which the mother cell traffics cytoplasmic SYS-1 to the centrosome, thus enriching SYS-1 fluorescence at the centrosome, while also internally controlling for bleaching via repetitive imaging. CEI measurements provide a clearer observation of the initial increase in the red channel of centrosomal DENDRA2::SYS-1 after photoconversion (Fig. 2G). CEI data show a similar rate of declining preexisting DENDRA2::SYS-1 centrosomal levels as the raw data but emphasizes the stability of newly translated centrosomal DENDRA2::SYS-1 enrichment, which is stable and low (Fig. 2G). Since we would expect no centrosomal enrichment of preexisting DENDRA2::SYS-1 if SYS-1 were trafficked to a different locale to be degraded, our finding that older SYS-1 preferentially localizes to the centrosomes is consistent with our model of SYS-1 enrichment from the cytoplasm and subsequent degradation at the centrosome. Altogether, these data suggest that newly synthesized and trafficked SYS-1, coupled with degradation of preexisting centrosomal SYS-1, lead to a stable steady state at the centrosomal locale, while indicating that the centrosome is the terminal locale of SYS-1. Notably, degradation is sequential; pre-existing centrosomal SYS-1 is degraded prior to newly trafficked SYS-1, which is stably replenished by cytoplasmic trafficking.

### Photoconversion of the cytoplasm of the P1 cell unveils dynamics of dynein trafficking of DENDRA2::SYS-1 during mitosis

Previous data indicate that microtubule trafficking via the dynein complex is needed for SYS-1 centrosomal localization [34]. By photoconverting subdomains of the posterior pole of the P1 cell cytoplasm, we tested the ability of the cell to traffic SYS-1 to the centrosomes. Following initial imaging during early mitosis, where pre-conversion levels of centrosomal DENDRA2::SYS-1 were measured with both green and red emission, a portion of the cytoplasm was photoconverted (Fig. 3A). Embryos were imaged every 20 seconds during mitosis (Fig. 3B) and DENDRA2::SYS-1 levels were measured in the proximal and the distal centrosomes in both the green and red channels. The photoconverted, red DENDRA2::SYS-1 represents newly trafficked SYS-1 from the anterior cytoplasmic region of the cell, while green DENDRA2::SYS-1 denotes older pre-existing centrosomal SYS-1 and newly trafficked SYS-1 from the non-photoconverted cytoplasmic SYS-1. Quantification of red newly trafficked SYS-1 showed centrosomal localization in both the proximal and distal centrosomes 20 seconds after photoconversion (Fig.3C-D), suggesting that SYS-1 is highly mobile and quickly trafficked to both centrosomes in early mitosis. However, though photoconverted SYS-1 accumulates on both proximal and distal centrosomes, the rates of accumulation are not equal. When directly comparing red newly trafficked DENDRA2::SYS-1 levels in the proximal and distal centrosomes, we observe that proximal centrosomes have a higher enrichment of SYS-1 throughout mitosis when compared to distal centrosomes (Fig. 3D). The difference in centrosomal DENDRA2::SYS-1 levels between the centrosomes suggest that cytoplasmic SYS-1 from any locale can be trafficked to both centrosomes but is more likely to be trapped and trafficked via dynein to the most proximal centrosome.

**Figure 3:**
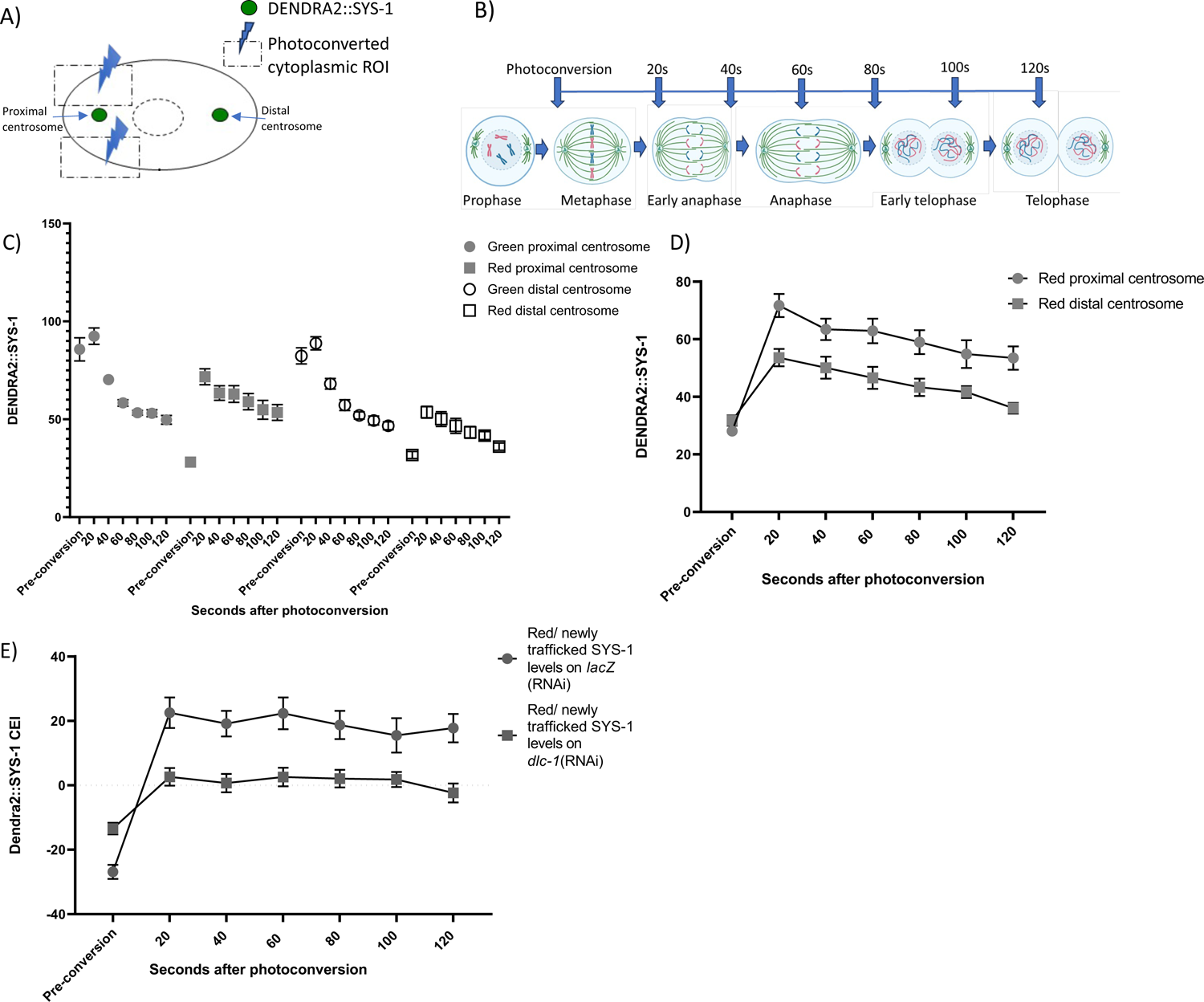
Photoconversion of the cytoplasm of the P1 cell unveils dynamics of dynein trafficking of DENDRA2::SYS-1 during mitosis. A) Diagram of photoconversion of the P1 cell. Dotted square represents the photoconverted area of the cytoplasm at metaphase. B) Photoconversion protocol for cytoplasm photoconversion in C), D), and E). C) Quantification of raw DENDRA2::SYS-1 of the distal and proximal centrosome in green and red channels pre- and post-photoconversion of the cytoplasm, measured over a 2 min. timecourse in 20 s intervals after photoconversion on control *lacZ*(RNAi). Mean +/−SEM. N=12. D) Quantification of DENDRA2::SYS-1 of the distal and proximal centrosome in red channel pre- and post-photoconversion of the cytoplasm, measured over a 2 min. timecourse in 20 s intervals after photoconversion on control *lacZ*(RNAi). Mean +/−SEM. N=12. E) Timecourse depicting red, photoconverted DENDRA2::SYS-1 CEI on *lacZ*(RNAi) and *dlc-1*(RNAi). Mean +/−SEM. N= 12 and 15, respectively.

To test the role of microtubule trafficking on DENDRA2::SYS-1, we depleted the light subunit of the dynein complex, DLC-1, via RNAi and measured the CEI of photoconverted DENDRA2::SYS-1 in the proximal centrosome. Previous results show DLC-1 is required for efficient centrosomal enrichment of SYS-1 [34]. After *dlc-1(RNAi),* the red, newly trafficked DENDRA2::SYS-1 is stable throughout mitosis; however, DLC-1 knockdown hinders SYS-1 trafficking to the centrosome, resulting in a decrease of centrosomal SYS-1 enrichment throughout mitosis compared to controls (Fig. 3E). This result is consistent with impaired SYS-1 centrosomal recruitment observed in cells lacking efficient dynein trafficking [34]. These data, which is consistent with our model of SYS-1 capture and centrosomal trafficking, suggest microtubule trafficking via the dynein complex is key for centrosomal SYS-1 localization and subsequent degradation.

### E and MS inheritance of mother cell SYS-1 is limited by EMS centrosomal processing

To determine the extent to which changes in SYS-1 centrosomal trafficking and degradation affect inheritance of mother cell SYS-1 into daughter cells, we photoconverted the entire EMS cell, the first ACD driven by the Wnt/β-catenin asymmetry pathway [39], of DENDRA-2::SYS-1-expressing embryos at cytokinesis and measured inherited (red) SYS-1 in the E and MS daughter cell nuclei. Here, E is the Wnt-signaled cell that normally exhibits higher SYS-1 levels and MS is the unsignaled cell with lower SYS-1 levels [21, 23]. For photoconversion of DENDRA2::SYS-1, we exposed EMS to UV (405nm) [53] at 100% laser power for 36 iterations during EMS cytokinesis. Subsequently, we waited two minutes for the nuclear envelope of the E and MS daughters to reform and imaged every 60 seconds for 7 minutes (Fig. 4A). Then, we normalized the nuclear fluorescence of DENDRA2::SYS-1 to the nuclear autofluorescence in wild-type (N2) embryos using identical measurements. Consistent with our FRAP studies, we observed inherited DENDRA2::SYS-1 present in both E and MS daughter cells (Fig. 4B). Interestingly, the amount of DENDRA2::SYS-1 inherited by the E daughter was higher than that present in MS (Fig. 4B), perhaps reflecting greater stability of inherited SYS-1 in the signaled E cell. We also noted that MS inheritance, while lower than E, was on average above autofluorescence. This result indicates SYS-1 inheritance persists even in the unsignaled MS cell. The timecourse also allowed us to observe changes in nuclear DENDRA2::SYS-1 over the cell cycle of the E and MS cells, up to 9 minutes after EMS division. For both the E and MS daughter cells, we found inherited nuclear DENDRA2::SYS-1 levels to remain stable throughout the timecourse (Fig. 4B), indicating the asymmetry in inherited nuclear DENDRA2::SYS-1 occurs shortly after EMS division.

**Figure 4:**
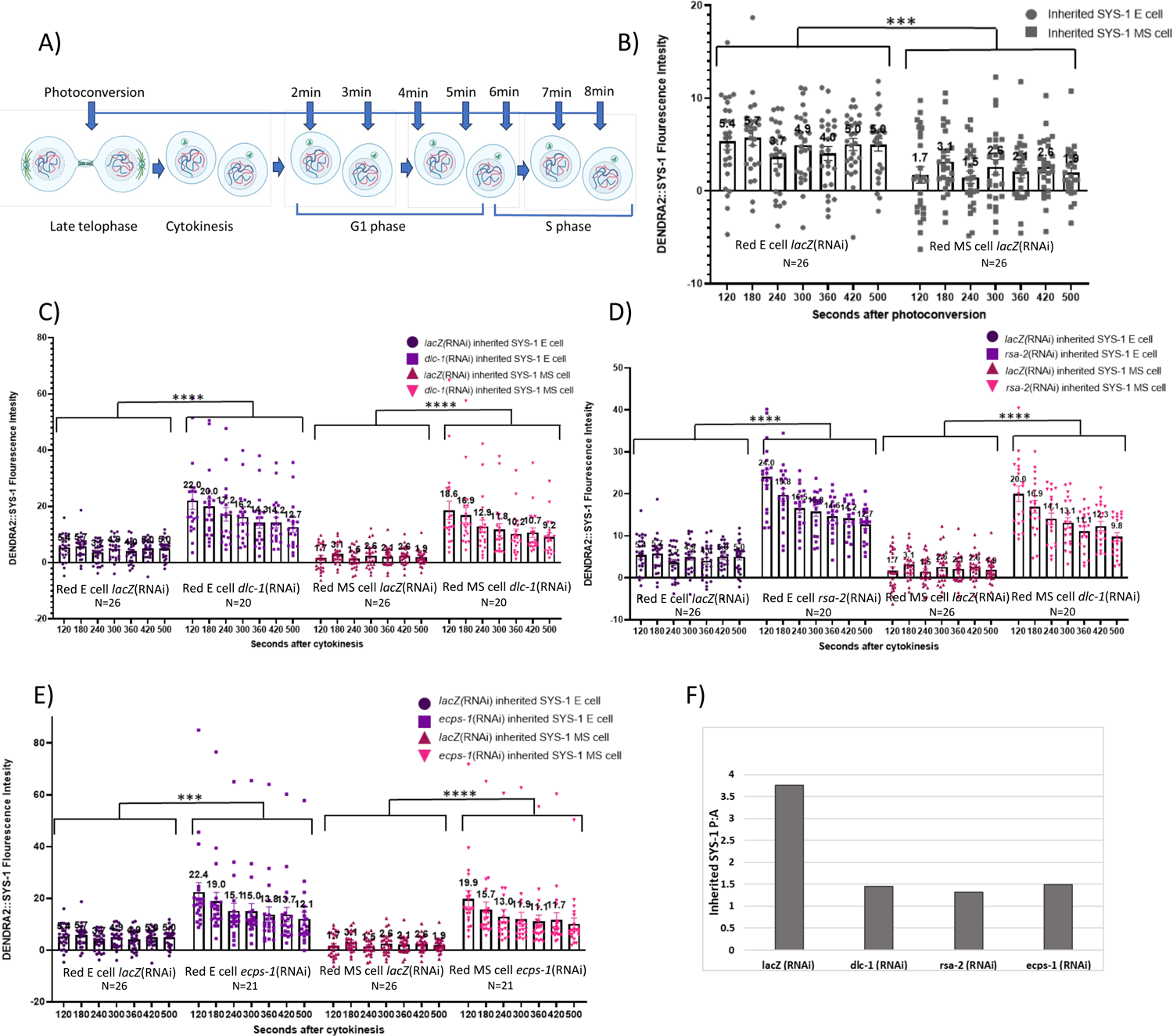
E and MS inheritance of mother cell SYS-1 is limited by EMS centrosomal processing. A) Photoconversion protocol for quantification of nuclear inherited DENDRA2::SYS-1 timecourse. B) Quantification of inherited, red DENDRA2::SYS-1 nuclear levels of the E and MS cells at different timepoints starting 2 m after cytokinesis of the EMS mother cell in 60 second increments on the control *lacZ*(RNAi). Mean +/−SEM. 2way ANOVA followed by Tukey Mixed effects analysis in Multiple Comparisons test. N=12. C) Quantification of inherited, red DENDRA2::SYS-1 nuclear levels of the E and MS cells timecourse on control vs. *dlc-1*(RNAi) (N=26 and N=20, respectively). Mean +/−SEM. 2way ANOVA followed by Sidak Mixed effects analysis in Multiple Comparisons test. D) Quantification of inherited, red DENDRA2::SYS-1 nuclear levels of the E and MS daughter cells timecourse on control vs. *rsa-2*(RNAi) (N=26 and N=20, respectively). Mean +/−SEM. 2way ANOVA followed by Sidak Mixed effects analysis in Multiple Comparisons test. E) Quantification of inherited, red DENDRA2::SYS-1 nuclear levels of the E and MS daughter cells timecourse on control vs. *ecps-1*(RNAi) (n=26 and n=20, respectively). Mean +/−SEM. 2way ANOVA followed by Sidak Mixed effects analysis in Multiple Comparisons test. F) Quantification of fold change in asymmetry between the E and MS cells in *lacZ, dlc-1, rsa-2 and ecps-1,* (RNAi). DENDRA2::SYS-1 fluorescent intensity was calculated by subtracting N2 fluorescence at all different timepoints. * represent the P-value when comparing the averaged value of all time points of *lacZ(*RNAi) to experimental condition. Table S1. P-values when comparing each time point. * p ≤ 0.05, ** p ≤ 0.01, *** p ≤ 0.001, **** p ≤ 0.0001

Next, we tested the requirement for dynein trafficking and centrosomal localization in SYS-1 inheritance. We predicted that impaired mother cell SYS-1 processing would lead to an increase in inherited SYS-1 levels, given the previously observed changes in cell fate under these conditions (DLC-1, RSA-2 and ECPS-1 knockdowns) [34]. We observed that when microtubule trafficking is impaired by the knockdown of DLC-1 there is a significant increase in the amount of inherited DENDRA2::SYS-1 nuclear levels of both the E and MS cells (Fig. 4C). Initial levels of inherited SYS-1 were 3 and 10-fold higher in the E and MS cells, respectively (Fig. 4C), suggesting mother cell dynein trafficking is required to limit the amount of inherited SYS-1.

We further investigated the role of SYS-1 centrosomal localization in controlling SYS-1 inheritance by preventing centrosomal localization of SYS-1 via knockdown of the centrosomal scaffold protein RSA-2. Loss of centrosomal localization by *rsa-2(RNAi)* increased the amount of inherited nuclear DENDRA2::SYS-1 by 3.4- and 11-fold in E and MS daughter cells, respectively (Fig. 4D). Lastly, we impaired the degradation of centrosomal SYS-1 by knocking down the adaptor protein ECPS-1, the worm homolog to EMC29, (a proteasome-dynein adaptor and scaffold) [35, 55]. Previous studies showed that ECPS-1 knockdown increases SYS-1 levels at the centrosome, suggesting a role promoting SYS-1 centrosomal degradation [34]. *ecps-1*(RNAi) lead to an increase of the amount of DENDRA2::SYS-1 inherited into the E and MS cells by 3- and 10.7-fold, respectively (Fig. 4E). These data suggest that the various mechanisms required for mother cell centrosomal SYS-1 degradation are normally required to prevent SYS-1 over-inheritance.

Finally, to assess the effect of the over-inherited SYS-1 on loss of daughter cell SYS-1 asymmetry and further loss of the asymmetric cell division of EMS, we averaged the inherited DENDRA2::SYS-1 nuclear levels in each condition throughout the 7-minute timecourse and measured the fold change between the E and the MS daughters. These data showed a loss of asymmetry of inherited SYS-1 in all the experimental conditions tested compared to control RNAi (Fig. 4F). The data suggest that SYS-1 is normally inherited into both the E and the MS daughter cells and that, early in the cell cycle, the asymmetry between E and MS is established. For proper inheritance of SYS-1 during the EMS ACD the mother cell requires SYS-1 centrosomal trafficking, coupling, and degradation (also referred to as centrosomal processing) to prevent over-inheritance of SYS-1 and loss of sister cell asymmetry.

### Loss of centrosomal SYS-1 results in increased levels of nuclear *de novo* SYS-1 in EMS daughter cells

Given the fact that centrosomal processing functions in the mother cell limit SYS-1 inheritance, we predicted that mother cell, but not *de novo* translated DENDRA2::SYS-1 would be affected when mother cell centrosomal processing of SYS-1 is impaired. To test this, we photoconverted at EMS cytokinesis and measured green/*de novo* DENDRA2::SYS-1 nuclear levels in E and MS daughter cells starting 2 minutes after cytokinesis and in 60 second increments for 7 minutes (Fig. 5A, B). We observed that both the E cell, the Wnt-signaled cell, and MS, the unsignaled cell, were actively translating DENDRA2::SYS-1 (Fig. 5A), consistent with our FRAP results (Fig.1C). The E cell also showed higher levels of *de novo* nuclear DENDRA2::SYS-1 levels than the MS cell (3.6-fold increase in E vs MS), indicating robust asymmetry in synthesis or stability of newly translated DENDRA2::SYS-1 (Fig. 5B). Additionally, the timecourse showed that the E and MS *de novo* nuclear DENDRA2::SYS-1 levels remained stable (Fig. 5B). These data suggest that SYS-1 synthesis occurs in both the signaled E and unsignaled MS daughter cells and that the rate of synthesis:degradation is higher in E than MS.

**Figure 5:**
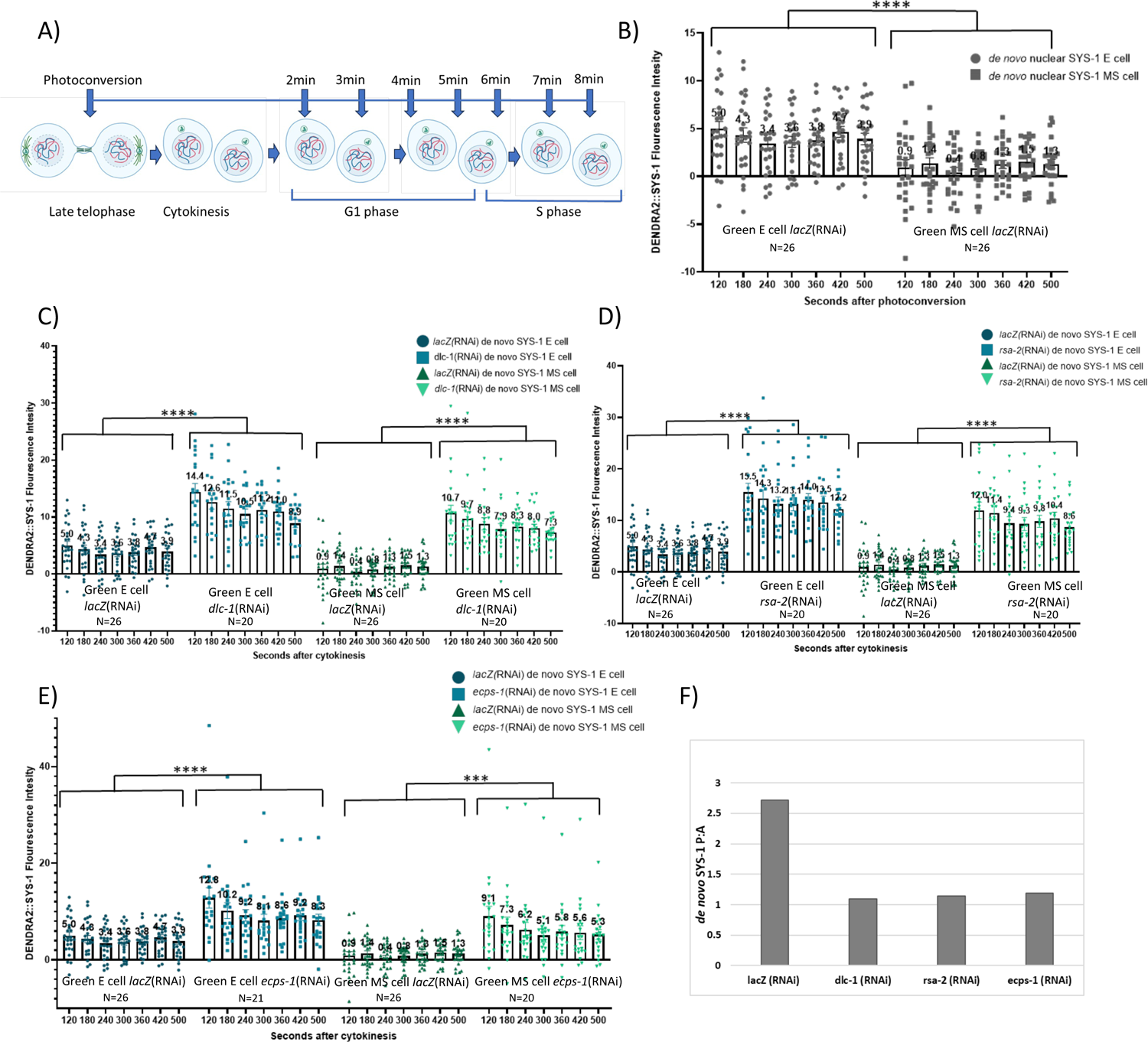
Loss of centrosomal SYS-1 results in increased levels of nuclear *de novo* SYS-1 in EMS daughter cells. A) Photoconversion protocol for quantification of nuclear inherited DENDRA2::SYS-1 timecourse. B) Quantification of green/*de novo* SYS-1 nuclear levels in the E and MS cells after photoconversion of the EMS cell at cytokinesis; the timecourse depicts SYS-1 nuclear levels at different timepoints starting 2 minutes after cytokinesis of the EMS cell in 60 second increments on the control *lacZ*(RNAi) (n=26). Mean+/− SEM. 2way ANOVA followed by Sidak Mixed effects analysis in Multiple Comparisons test. C) Quantification of *de novo* SYS-1 nuclear levels of the E and MS cells timecourse on *lacZ(RNAi)* and *dlc-1*(RNAi) (n=26 and n=20, respectively). Mean +/−SEM. 2way ANOVA followed by Sidak Mixed effects analysis in Multiple Comparisons test. D) Quantification of *de novo* SYS-1 nuclear levels of the E and MS cells timecourse on *lacZ(RNAi)* and *rsa-2*(RNAi) (n=26 and n=20, respectively). Mean +/−SEM. 2way ANOVA followed by Sidak Mixed effects analysis in Multiple Comparisons test. E) Quantification of *de novo* SYS-1 nuclear levels of the E and MS cells timecourse on *lacZ(RNAi)* and *ecps-1*(RNAi) (n=26 and n=20, respectively). Mean+/− SEM. 2way ANOVA followed by Sidak Mixed effects analysis in Multiple Comparisons test. F) Quantification of fold change in asymmetry between the E and MS daughter cell in *lacZ, dlc-1, rsa-2 and ecps-1* (RNAi). DENDRA2::SYS-1 fluorescent intensity was calculated by subtracting N2 fluorescence at all different timepoints. * represent the P-value when comparing all time points and the average of control to experimental condition. Table S1. P-values when comparing each time point. * p ≤ 0.05, ** p ≤ 0.01, *** p ≤ 0.001, **** p ≤ 0.0001

Subsequently, we observed *de novo* nuclear DENDRA2::SYS-1 levels in the daughter cells after knocking down DLC-1, therefore impairing microtubule trafficking of SYS-1 to the centrosome in the mother cell [34]. This surprisingly resulted in a significant increase in the amount of *de novo* nuclear SYS-1 levels in both daughter cells at all time points (Fig. 5C). This increase in *de novo* SYS-1 was also observed in the other tested conditions that showed increased DENDRA2::SYS-1 inheritance, *rsa-2* and *ecps-1 (*RNAi) (Fig. 5D,E). In all the tested conditions (*dlc-1, rsa-2* and *ecps-1 (*RNAi)), we also observed a loss of asymmetry compared to the control (Fig. 5F).

Thus, we find that impairing proper SYS-1 centrosomal processing during mitosis of the EMS cell showed increased levels of both inherited and *de novo* nuclear DENDRA2::SYS-1 in EMS daughter cells (Figs. 4 and 5). Additionally, we observed that over time in the tested conditions, nuclear DENDRA2::SYS-1 consistently decreases in backgrounds with higher than normal SYS-1 compared to wild type, which stably maintains its SYS-1 levels in E and MS (Table 1). This pattern of increased rate of DENDRA2::SYS-1 clearance was observed in both DENDRA2::SYS-1 pools, suggesting that both *de novo* and inherited SYS-1 are processed similarly in both daughter cells. These data suggest that over-inheritance of SYS-1 during asymmetric cell division affects the *de novo* SYS-1 that is generated or maintained in both daughter cells. This led us to hypothesize that the β-catenin “destruction complex” activity in daughter cells has a limited capacity and is not fully able to regulate SYS-1 in situations with increased levels of inherited SYS-1, an idea we test below.

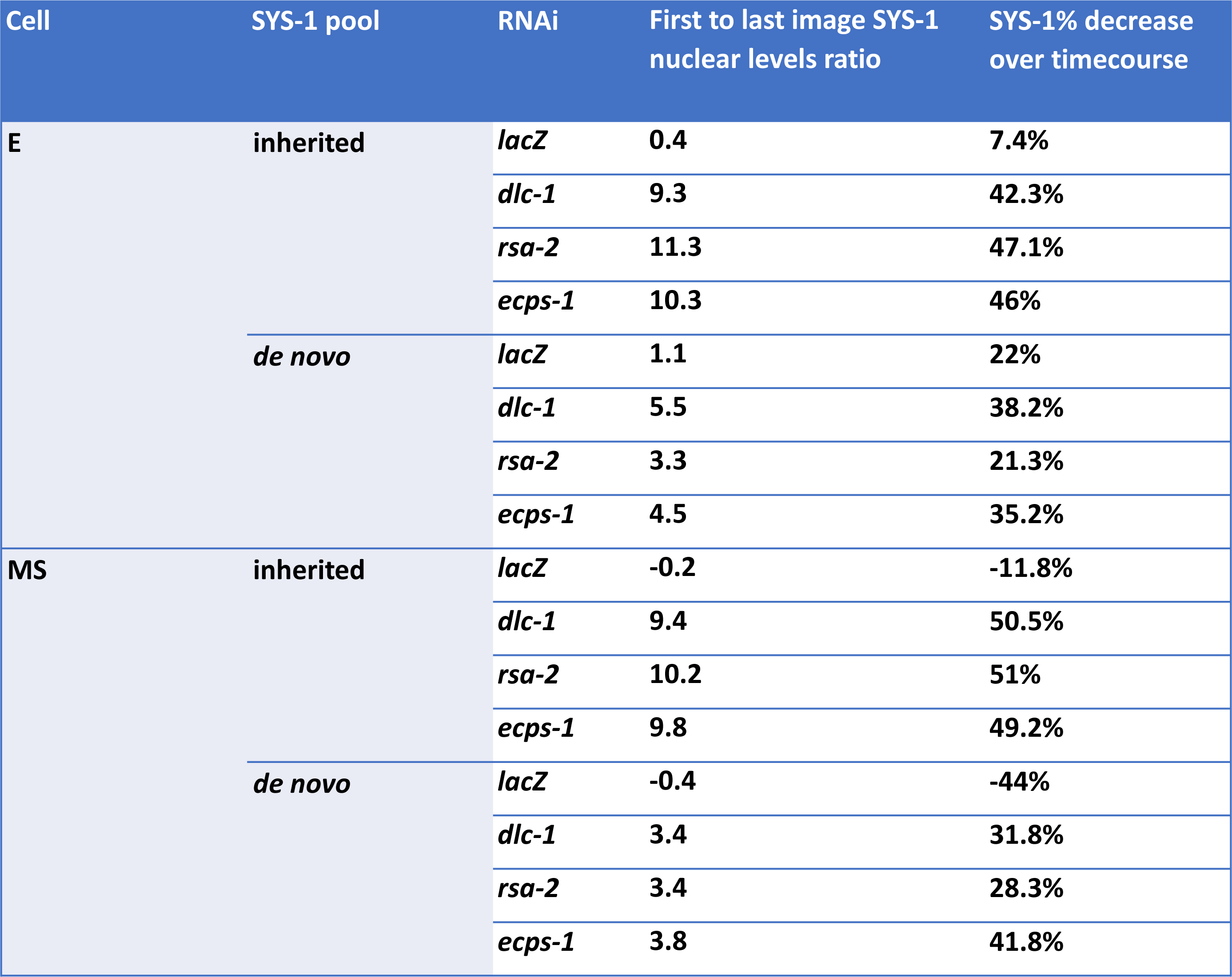

### Partial loss of the negative regulator of the Wnt signaling pathway, APR-1, affects both pools of nuclear SYS-1 in the E and MS cells

We targeted APR-1 a negative regulator of SYS-1 in the MS cell and elsewhere [21, 29] and a homolog of vertebrate APC, which is part of the β-catenin destruction complex and functions to limit β-catenin cytoplasmic accumulation in the Wnt-unsignaled state. APC also limits Wnt pathway activity after initial activation by targeting β-catenin for proteasomal degradation [56]. As expected, we found that depletion of APR-1 via RNAi led to a significant increase in *de novo* DENDRA2::SYS-1 nuclear levels throughout the MS cell cycle (Fig. 6A). We also observed a 2-fold increase in *de novo* DENDRA2::SYS-1 during the E cell cycle (Fig. 6A), which has not been shown previously. This result was unexpected as a role of APR-1 as a negative regulator of SYS-1 had been primarily observed in the unsignaled MS cell, which displays higher APR-1 levels [21, 26, 29, 57], while these data suggest APR-1 regulates SYS-1 levels in both daughter cells.

**Figure 6:**
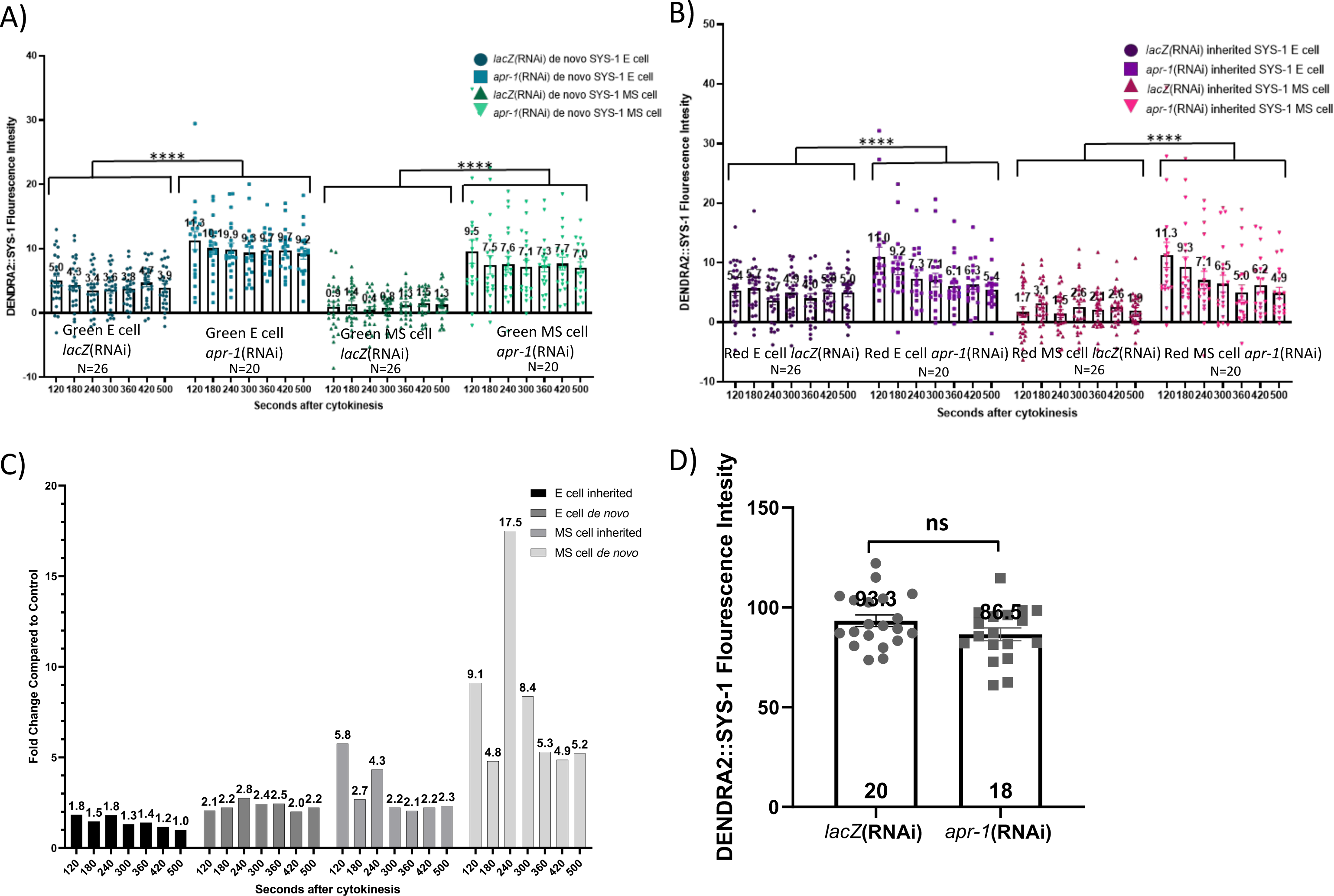
Partial loss of the negative regulator of the Wnt signaling pathway, APR-1, affects both pools of nuclear SYS-1 in the E and MS cells. A) Quantification of *de novo* SYS-1 nuclear levels in the E and MS cells after photoconversion at cytokinesis over 7-minute timecourse on *lacZ(RNAi) and apr-1* (RNAi) (n=26 and n=20, respectively). Mean +/−SEM. 2way ANOVA followed by Sidak Mixed effects analysis in Multiple Comparisons test. B) Quantification of inherited SYS-1 nuclear levels in the E and MS cells after photoconversion at cytokinesis over 7-minute timecourse on *lacZ(RNAi) and apr-1* (RNAi). Mean +/−SEM. 2way ANOVA followed by Sidak Mixed effects analysis in Multiple Comparisons test. C) Graph representation of the fold change in *de* novo and inherited nuclear SYS-1 of APR-1 compared to the control in the E and MS cells. D) Quantification of centrosomal DENDRA2::SYS-1 levels during telophase in the EMS cell on *lacZ(RNAi)* and *apr-1* (RNAi) (n=20 and n=18, respectively). Mean +/−SEM. Unpaired two-tail t-test. * represent the P-value when comparing all time points and the average of control to experimental condition. Table S1. P-values when comparing each time point. DENDRA2::SYS-1 fluorescent intensity was calculated by subtracting N2 fluorescence at all different timepoints. * p ≤ 0.05, ** p ≤ 0.01, *** p ≤ 0.001, **** p ≤ 0.0001

Additionally, we observe different nuclear DENDRA2::SYS-1 dynamics when APR-1 is knocked down compared to other backgrounds with elevated *de novo* DENDRA2::SYS-1. The E and MS cells *de novo* DENDRA2::SYS-1 levels show a smaller decrease compared to the DLC-1, RSA-2, and ECPS-1 knockdowns (Table 1), over the 7-minute timecourse (Fig. 6A). These results suggest that *de novo* SYS-1 in both daughter cells is limited by the destruction complex in wild type and in situations with elevated inherited SYS-1.

We also observed increased levels of inherited nuclear SYS-1 in both E and MS daughter cells after APR-1 depletion (Fig. 6B). Though the effect was visible in E, the highest fold change of *de novo* nuclear DENDRA2::SYS-1 after APR-1 depletion was in MS (Fig. 6C). In MS, the *de novo* pool was more highly affected by loss of APR-1 than the inherited pool, suggesting that, in wild type, APR-1 is more critical for negatively regulation of *de novo* SYS-1 (Fig. 6C). Though the effect of APR-1 depletion on inherited SYS-1 in MS leads to an average of 3.1-fold increase compared to wild type; this was less than the 7.9-fold increase we see in the MS *de novo* SYS-1 pool. By contrast, the difference between E inherited and E *de novo* SYS-1 after APR-1 depletion is not as striking (1.4-fold and 2.3-fold increase compared to wild type, respectively) (Fig. 6C). Therefore, the major regulatory function of APR-1 on SYS-1 nuclear levels in EMS ACD is to limit accumulation of *de novo* SYS-1 in MS. In the E cell, APR-1 plays a minor role in SYS-1 regulation, likely due to APR-1 repression due to Wnt signaling.

Despite the decreased effect of APR-1 depletion on DENDRA2::SYS-1 nuclear levels in E, our results suggest that APR-1 is also functioning in this Wnt-signaled daughter cell. However, an alternative hypothesis explaining our increased SYS-1 levels in E after APR-1 depletion is that APR-1 negatively regulates SYS-1 earlier in the EMS lineage. In this case, increased SYS-1 inheritance is would be expected to increase the total SYS-1 levels and possibly the stability of *de novo* SYS-1 in the daughter cells. To distinguish between these possibilities, we knocked down APR-1 and observed changes in total centrosomal DENDRA2::SYS-1 levels in the EMS cell during mitosis with no photoconversion. Centrosomal SYS-1 quantitation is a good readout of SYS-1 levels in the EMS cell and its progenitors because SYS-1 centrosomal accumulation reflects cytoplasmic SYS-1 levels [30] and any effect of SYS-1 levels can be clearly observed immediately before the cell divides, providing the latest timepoint possible before the birth of E and MS. We found that there is no significant change in centrosomal DENDRA2::SYS-1 levels during telophase of the EMS cell on *apr-1(RNAi)* compared to wild type (Fig. 6D), suggesting SYS-1 is not negatively regulated by APR-1 in EMS or its progenitors. Thus, our observations suggest that when the inherited SYS-1 pool is affected in the EMS mother cell, the *de novo* SYS-1 pools are secondarily affected in the daughter cells. Additionally, here we report a role for APR-1 limiting the accumulation of both SYS-1 pools in both the signaled and unsignaled EMS daughter cells.

## Discussion

In *C. elegans,* successful asymmetric cell divisions throughout development depend on proper regulation of SYS-1 levels. Here we show that in wild-type conditions, proper SYS-1 centrosomal degradation in the EMS mother cell is key to regulate inheritance into the E and MS cells during ACD. It was previously established that SYS-1 centrosomal trafficking, coupling and degradation in the mother cell is key for regulation of limiting subsequent daughter cell SYS-1 levels and subsequent asymmetric cell fate specification [30, 34]. These data suggested SYS-1 inheritance was normally limited by centrosomal degradation and that *de novo* synthesized SYS-1, regulated by asymmetric Wnt pathway activity, was the source of nuclear SYS-1 important for ACD. However, SYS-1 inheritance (or the lack thereof) and the role of centrosomal degradation have not been demonstrated. Photobleaching experiments show that mother cell bleaching of GFP::SYS-1 decreases the total GFP::SYS-1 in the daughter cells, suggesting that inherited SYS-1 was present in daughter cell nuclei in wild-type conditions. DENDRA2::SYS-1 photoconversion at cytokinesis directly confirmed DENDRA2::SYS-1 inheritance to both the E and the MS cell and that loss of centrosomal SYS-1 degradation not only increases the inherited DENDRA2::SYS-1 pool but also unexpectedly increases/stabilizes the *de novo* synthesized pool. We show that APR-1 regulates both inherited and *de novo* SYS-1 in E and MS, and the rate of clearance of extra inherited SYS-1 is dependent on APR-1 function. Photoconvertible SYS-1 also allowed us to test the model of SYS-1 trafficking to its final destination in the cell, the mitotic centrosome. Photoconversion of DENDRA2::SYS-1 at the centrosome during metaphase demonstrated that newly synthesized DENDRA2::SYS-1 is continually trafficked to the centrosome during mitosis, resulting in stable steady state SYS-1 levels despite this localization leading to increased SYS-1 turnover [30]. In contrast, preexisting SYS-1 is more enriched at the centrosome and decreases through mitosis. Thus, proper SYS-1 levels are dependent on both trafficking and degradation of SYS-1 in the mother cell along with negative regulation via the destruction complex in the daughter cells.

Based on our findings, we propose the following model (Fig. 7). In wild-type conditions during ACD of the EMS cell, a portion of cytoplasmic SYS-1 is trafficked to the centrosome where the scaffold protein RSA-2 localizes SYS-1 to the centrosome. Centrosomal SYS-1 is degraded by the centrosomal proteosome, while the remaining cytoplasmic SYS-1 is inherited into the E and MS daughter cells. Inherited SYS-1 is then differentially regulated by APR-1, primarily (but not exclusively) in MS cell. *De novo* SYS-1 also accumulates in both daughter cells, with the E cell having higher nuclear SYS-1 levels than the MS cell, and the *de novo* pool is similarly regulated by APR-1 as the inherited pool (Fig. 7A).

**Figure 7:**
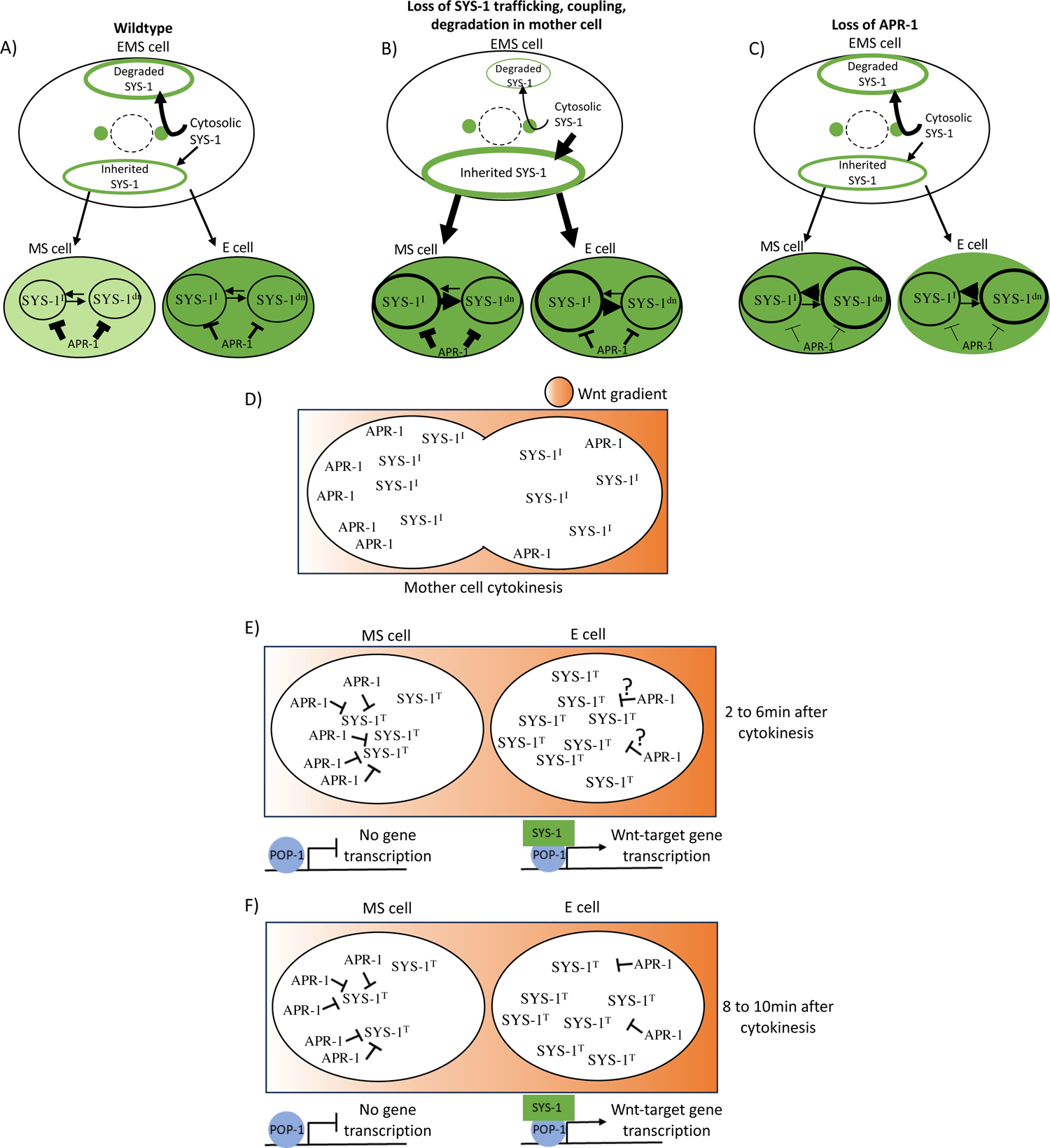
Model for SYS-1 regulation during ACD. See text for details. Weight of the arrows and lines represent the relative changed effect, oval size represents size of the relevant SYS-1 pool. SYS-1^i^, inherited SYS-1; SYS-1^dn^, *de novo* SYS-1.

Loss of mother cell regulation of centrosomal SYS-1 levels via depletion of RSA-2, DLC-1 or ECPS-1 prevents proper degradation of mother cell SYS-1 leading to over-inheritance of SYS-1 in both daughter cells. A higher rate of SYS-1 inheritance is correlated with higher *de novo* SYS-1 levels in both daughter cells, indicating an inability of the destruction complex to effectively regulate both SYS-1 pools due to dilution of the destruction complex activity. The overwhelming of the destruction complex allows SYS-1 nuclear levels to be higher in both the E and MS cell (Fig. 7B).

Loss of the negative regulator APR-1 has a direct effect in the *de novo* levels in both the E and the MS cells as we see no effect on EMS mother cell SYS-1 levels (Fig. 6D). Due to the similar regulation of *de novo* SYS-1 and inherited SYS-1 pools by APR-1, increased levels of both pools were observed after APR-1 depletion, however this was likely due to increased stability of just the *de novo* pool. A loss of APR-1 therefore leads to increased total SYS-1 levels and a loss of daughter cell asymmetry (Fig. 7C) [21].

Focusing on the EMS cell during cytokinesis, our model proposes that cell polarization by the Wnt gradient leads to differential distribution of APR-1. In the posterior region of the mother cell, APR-1 will not accumulate due to increased Wnt signaling (Fig. 7D). In the anterior region of the cell, as documented previously [58], there will be a greater accumulation of APR-1. This pattern of APR-1 asymmetry will drive asymmetric stability of SYS-1 post division (Fig. 7E,F). In the first portion of the daughter cell cycle, from two to six minutes post-cytokinesis (Fig 7E), the E cell increases its total nuclear SYS-1 levels, which encompasses the inherited and the *de novo* SYS-1 pools, due to decreased APR-1 levels and activity resulting from active Wnt signaling. During this same period, APR-1 in the MS cell decreases SYS-1 stability, leading to low total nuclear SYS-1 levels. Late during the cell cycle of the daughter cells, eight to ten minutes after cytokinesis (Fig. 7F), we observe decreased total SYS-1 levels in the E cell due to increasing negative regulation via APR-1. In the MS cell, a more pronounced SYS-1 decrease is observed, suggesting a lack of Wnt signaling and the negative regulation of *de novo* and inherited SYS-1 via APR-1 (Fig. 7F).

### Why is inherited SYS-1 asymmetric between E and MS?

Interestingly, the inherited DENDRA2::SYS-1 is not equal for both daughter cells; E shows higher levels of SYS-1 through the timecourse than its sister MS (Fig. 4A). Previous studies show a strong asymmetry of APR-1 during the EMS cell division, accumulating in the anterior side [26, 58]. Our data show that APR-1 predominantly functions in the MS cell, though we do also see APR-1 depletion effects in E (Fig. 6C). This suggests that SYS-1 inheritance may be symmetric [30], or at least correlates with the volume of the cytoplasm, but the stability of SYS-1 is asymmetric in the two daughter cells. In this model, inherited DENDRA2::SYS-1 in wild type is more stable and prevalent in the E cell (Fig. 4B). However, it remains feasible that asymmetric inheritance of other Wnt signaling components, such as Frizzled or Dishevelled, contribute to the asymmetry observed. Indeed, asymmetric MOM-5/Frizzled has been reported in the early embryonic blastomeres [37] and the seam cells [28], suggesting these components could be influencing the distribution of other Wnt signaling components during ACD. Together, these data suggest that while SYS-1 might be symmetrically inherited into the daughter cells, further SYS-1 regulation and stability is dependent on the asymmetric inheritance of positive and negative regulators of Wnt-signaling.

### Why does over-inheritance of SYS-1 correlate with higher *de novo* SYS-1 levels?

While loss of microtubule trafficking, centrosomal coupling or degradation of SYS-1 in the mother cell led to an increase in the amount of inherited DENDRA2::SYS-1, we also observed increased levels of *de novo* DENDRA-2::SYS-1 in both daughter cells. These unexpected results led to further investigation of the role of negative regulators of SYS-1 in the Wnt-signaling pathway. Since APR-1 depletion primarily affects *de novo* DENDRA2::SYS-1 rather than DENDRA2::SYS-1 levels in the mother cell (Fig 6A, D), we used this background to test the effect of increased *de novo* DENDRA2::SYS-1 on maintenance of the inherited DENDRA2::SYS-1 pool. We observed increased inherited nuclear DENDRA2::SYS-1 levels in both daughter cells (Fig. 6B). These results supported our hypothesis that when either of the pools is affected, this will influence the amount of nuclear SYS-1 in the other pool. These results could be explained by an overwhelming of the destruction complex that is equally targeting both *de novo* and inherited SYS-1 pools, thus leading to an accumulation of *de novo* SYS-1 as destruction complex activity is diluted by increased inherited SYS-1. Our data suggest that increased levels of SYS-1 in the daughter cells directly increases SYS-1 stability in the primarily affected pool (inherited DENDRA2::SYS-1 in *dlc-1, rsa-2 and ecps-1*(RNAi) or *de novo* DENDRA2::SYS-1 in *apr-1*(RNAi)), which in turn perturbs negative regulation of the other pool, leading to an increase in the total levels of nuclear SYS-1. Thus, we conclude that the SYS-1 pools are interchangeable, and both regulated by the destruction complex.

We also allowed tested the hypothesis that the rate of nuclear SYS-1 loss observed in cases of elevated inherited DENDRA2::SYS-1 levels (RSA-2, ECPS-1, or DLC-1 depletion) are due to the function of the destruction complex decreasing SYS-1 back to basal levels. APR-1 knockdown indeed led to a more moderate rate of decrease in *de novo* SYS-1 levels throughout the timecourse compared to the other conditions tested (*dlc-1, rsa-2 and ecps-1(RNAi)*), suggesting efficient clearance of extra SYS-1 requires APR-1 (Table1) (Fig. 6A).

### APR-1 functions in the E cell?

Our data suggest that the destruction complex functions in the Wnt-signaled E cell. When knocking down APR-1, we found increased levels of *de novo* nuclear DENDRA2::SYS-1 in both the E and the MS, and while this result was expected for the MS cell, we did not expect to see significant changes in the E cell (Fig. 6A and B). Previous studies investigated the kinetic responses of β-catenin to Wnt-signaling in mammalian cells and found that the destruction complex functions in both the absence and presence of Wnt-signaling. However, when Wnt ligand is present, the destruction complex is only partially active [59]. While this has not yet been described in *C. elegans*, our results support those findings. FRAP experiments demonstrated that *de novo* GFP::SYS-1 plateaus 6 to 8 minutes after cytokinesis, suggesting its stabilization is temporally limited in E (Fig. 1D), and this could be due to destruction complex function. Additionally, the partial depletion of APR-1 via RNAi in our experiments seems to lower the steady state of degradation when Wnt-signaling is present, which supports the observed role for the destruction complex in the E cell.

While our studies focused on the EMS daughter cells, previous studies showed that in both the daughters of E (Ea and Ep) and MS (MSa and MSp), SYS-1 asymmetry is lost after *apr-1*(RNAi) [21]. These results further support the idea that APR-1 regulation is key to maintain daughter cell asymmetry and its role in both E, the Wnt-signaled daughter cell and MS, the unsignaled cell. In addition, it has been previously shown that the observed asymmetry is not due to nuclear export or differential subcellular distribution of SYS-1, but to proteasomal degradation [21]. These data suggest that SYS-1 asymmetry is tightly regulated by proteasomal degradation via mother cell centrosomal processing and the destruction complex function in the EMS daughters.

Our proposed mechanism of SYS-1 regulation and inheritance during ACD sheds light on the tight regulation of ACD processes and the several layers of negative regulation needed for proper asymmetric target gene activation. Indeed, trafficking and degradation of mother cell SYS-1 maintains SYS-1 levels in the daughters at a level that can be effectively regulated by the destruction complex.

## Acknowledgments

We thank members of the Phillips lab for comments on the manuscript. Strains were provided by the Caenorhabditis Genetics Center, which is funded by the National Institutes of Health (NIH) Office of Research Infrastructure Programs (P40 OD01440). We also want to thank the Carver Center for Imaging at the University of Iowa and its director, Dr. Michael Dailey. This work was supported by NIH award RO1GM114007 (to B.T.P), and University of Iowa Graduate College Fellowships to M.V.

## Notes

### Competing Interest Statement

The authors have declared no competing interest.

### Summary of Updates

This version shortens and simplifies the text and includes a new data table.

