## Supplemental figures for "SYS-1/beta-catenin inheritance and regulation by Wnt-signaling during asymmetric cell division"

### Supplemental figure 1

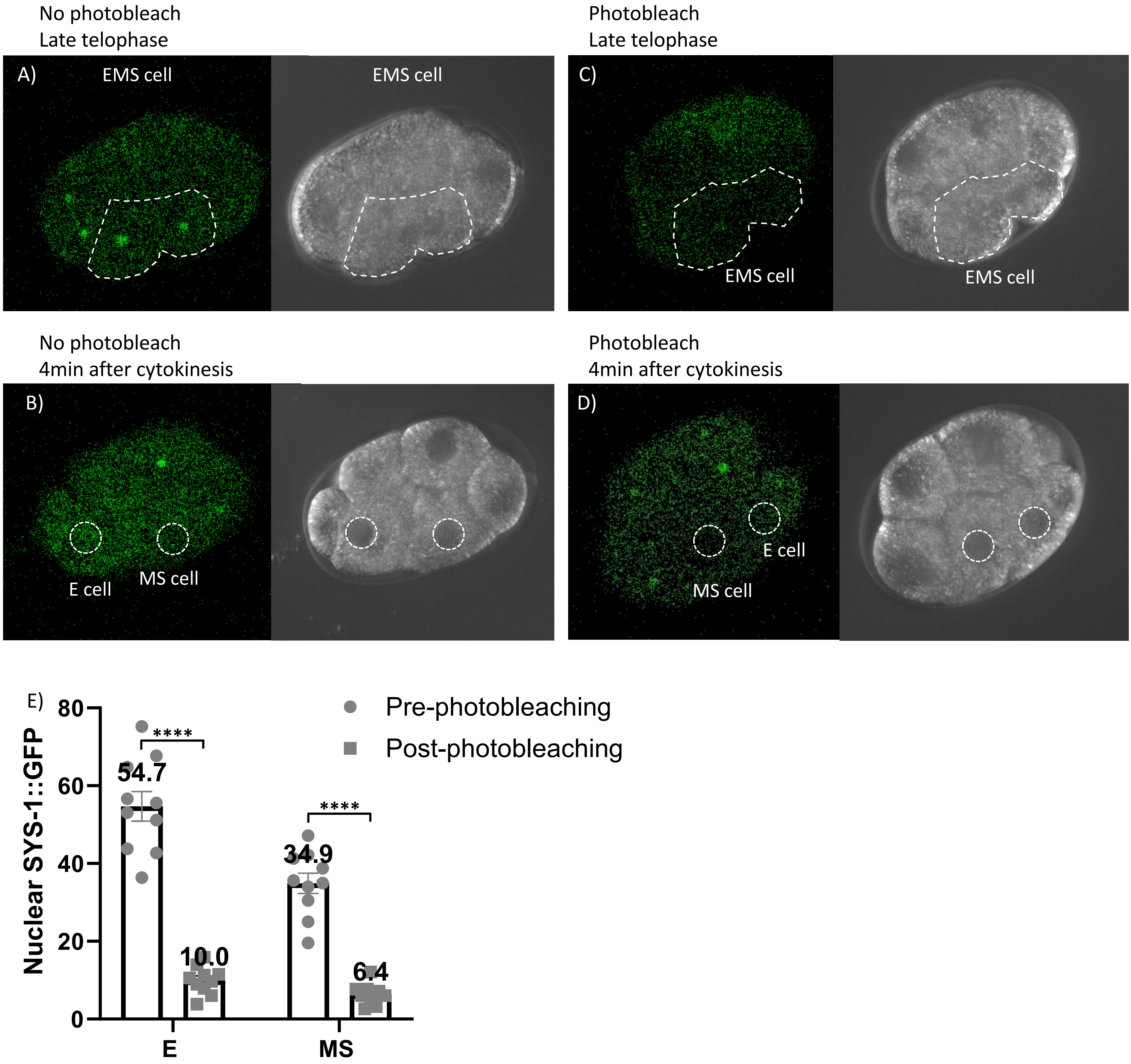

**Supplemental figure 1:**

**Photobleaching efficacy of SYS-1::GFP.** A)-D) Representative images of photobleaching experiments. A) No photobleach image of EMS cell taken late telophase (at time of photobleaching). B) No photobleach image of E and MS cell 4min after cytokinesis. C) EMS cell photobleached at late telophase. D) E an MS cells after being photobleached and imaged 4min after cytokinesis. D) Quantification of normalized SYS-1 to assess photobleaching efficacy, photobleach and imaged 3min after cytokinesis (n=10). SYS-1::GFP fluorescent intensity was calculated by subtracting N2 fluorescence. Mean +/-SEM. Paired two-tail t-test* $p \leq 0.05$, ** $p \leq 0.01$, *** $p \leq 0.001$, **** $p \leq 0.0001$

### Supplemental figure 2

A)

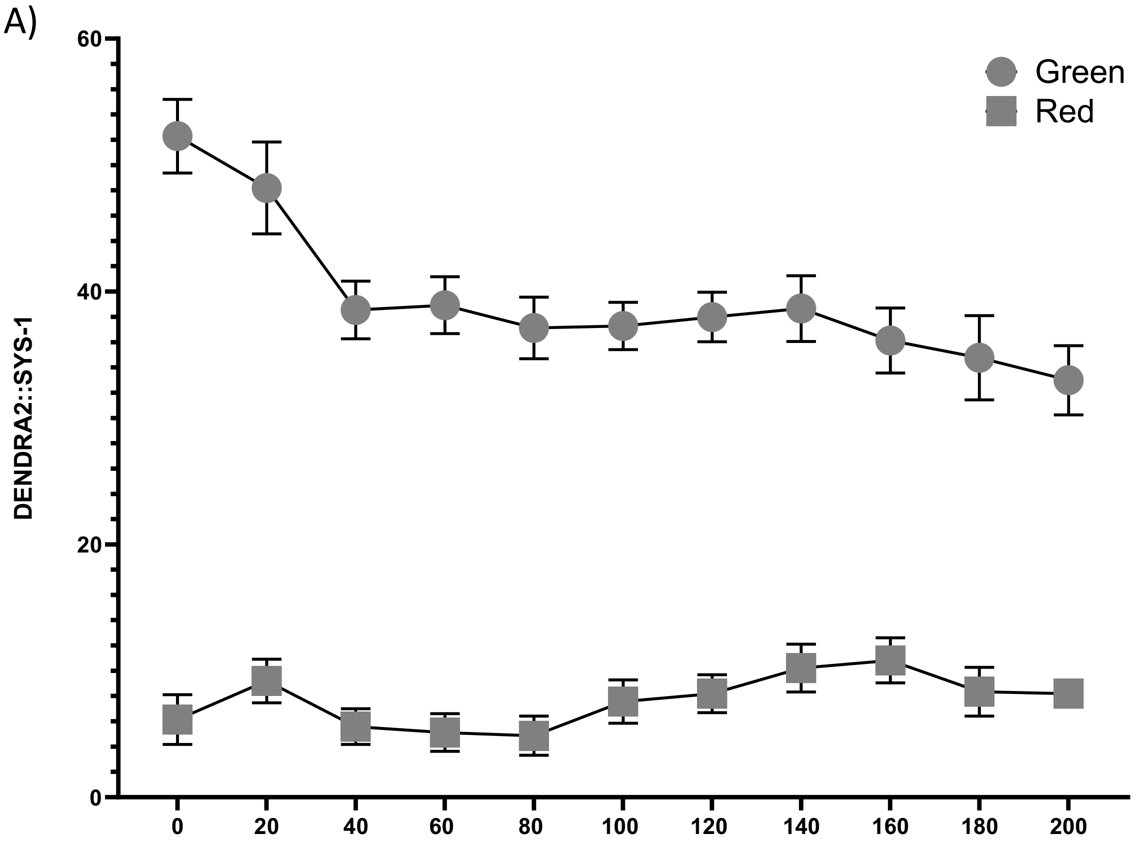


B)

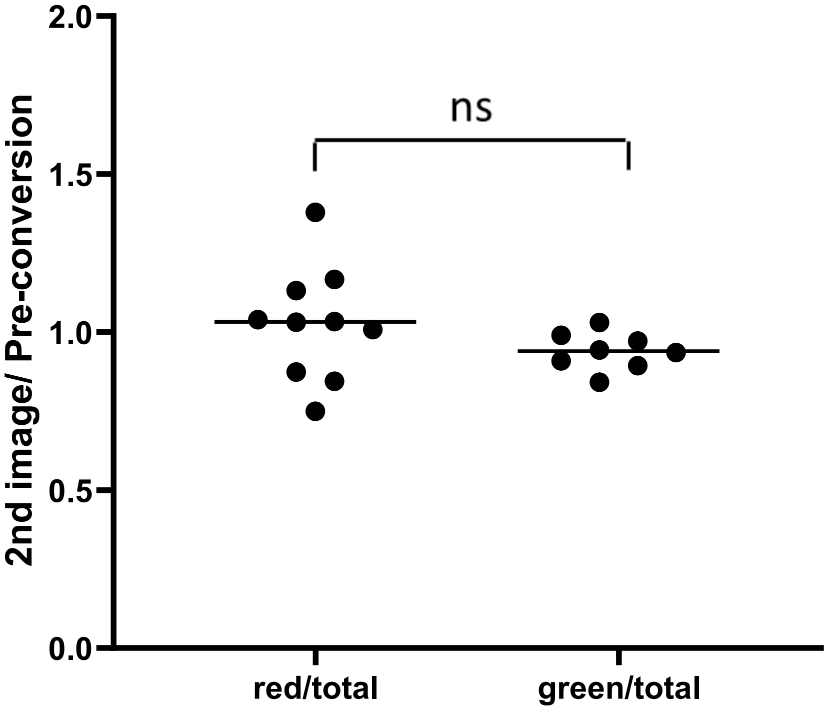

**Supplemental figure 2:**

**Photoconversion efficacy of DENDRA2::SYS-1.** A) Quantification of average centrosomal DENDRA2::SYS-1 in both red and green channels without photoconversion normalized to N2, measured over a 3.5 min. timecourse in 20 s intervals on control *lacZ*(RNAi). Mean +/- SEM. N=7. B) Average of ratios to assess photoconversion efficacy, red/total ratio represent DENDRA2::SYS-1 centrosomal red levels immediately after photoconversion divided by green pre-conversion levels (N=10). Green/total ratio represent DENDRA2::SYS-1 centrosomal green levels after fist image was taken divided by total green pre-conversion levels (N=8). DENDRA2::SYS-1 fluorescent intensity was calculated by subtracting N2 fluorescence. Mean +/-SEM. Paired two-tail t-test* $p \leq 0.05$, ** $p \leq 0.01$, *** $p \leq 0.001$, **** $p \leq 0.0001$
